# Investigating the roles of *Listeria monocytogenes* peroxidases in growth and virulence

**DOI:** 10.1101/2021.04.12.439369

**Authors:** Monica R. Cesinger, Nicole H. Schwardt, Cortney R. Halsey, Maureen K. Thomason, Michelle L. Reniere

## Abstract

Bacteria have necessarily evolved a protective arsenal of proteins to contend with peroxides and other reactive oxygen species generated in aerobic environments. *Listeria monocytogenes* encounters an onslaught of peroxide both in the environment and during infection of the mammalian host, where it is the causative agent of the foodborne illness listeriosis. Despite the importance of peroxide for the immune response to bacterial infection, the strategy by which *L. monocytogenes* protects against peroxide toxicity has yet to be illuminated. Here, we investigated the expression and essentiality of all the peroxidase-encoding genes during *L. monocytogenes* growth *in vitro* and during infection of murine cells in tissue culture. We found that *chdC* and *kat* were required for aerobic growth *in vitro*, and *fri* and *ahpA* were each required for *L. monocytogenes* to survive acute peroxide stress. Despite increased expression of *fri, ahpA*, and *kat* during infection of macrophages, only *fri* proved necessary for cytosolic growth and intercellular spread. In contrast, the proteins encoded by *lmo0367, lmo0983, tpx, lmo1609*, and *ohrA* were dispensable for aerobic growth, acute peroxide detoxification, and infection. Together, our results provide insight into the multifaceted *L. monocytogenes* peroxide detoxification strategy and demonstrate that *L. monocytogenes* encodes a functionally diverse set of peroxidase enzymes.

## INTRODUCTION

Bacteria that replicate in aerobic environments must contend with reactive oxygen species (ROS) such as superoxide O_2_^-^), hydroxyl radicals (HO•), and hydrogen peroxide (H_2_O_2_). These ROS result from the incomplete reduction of molecular oxygen and are both produced by the bacteria (endogenously) and encountered extracellularly (exogenous). Accumulation of ROS leads to DNA damage and mutagenesis, protein oxidation, and destruction of iron-sulfur clusters and other iron-containing proteins (1). In light of these deleterious effects, it is perhaps unsurprising that the mammalian immune system uses ROS to defend against invading pathogens via the phagocytic respiratory burst, a period of increased oxygen consumption observed during phagocytosis due to the activity of NADPH oxidase (2). NADPH oxidase generates ROS in the phagosome and is critical to the innate immune response, as evidenced by the increased susceptibility of patients with chronic granulomatous disease (CGD) to severe infections. CGD is an inherited immunodeficiency caused by deletions or mutations in the genes encoding NADPH oxidase (3). Without this immune defense mechanism, CGD patients suffer from invasive and recurrent infections (4).

Bacteria employ many defense mechanisms to defend themselves against ROS-mediated oxidative stress, the most impactful being the production of scavenging enzymes (5). Superoxide dismutase detoxifies superoxide, generating molecular oxygen and hydrogen peroxide, which is then efficiently scavenged by peroxidases and catalases. Peroxidases are enzymes that reduce hydrogen peroxide and are the primary scavengers at low concentrations of peroxide. At high concentrations of peroxide when peroxidases become saturated, catalases disproportionate peroxide and become the dominant scavengers (6). Bacteria typically produce multiple functionally redundant enzymes to protect against peroxides. In fact, peroxide toxicity is not detected in the model organism *Escherichia coli* unless three peroxide scavenging enzymes are deleted simultaneously (*ahpCF, katG*, and *katE*) (7).

While the mechanisms of peroxide detoxification have been studied for decades in the model organisms *E. coli* and *Bacillus subtilis* (5, 6, 8), the mechanisms by which *Listeria monocytogenes* defends against ROS are less clear. *L. monocytogenes* is a facultative foodborne pathogen that invades host cells, replicates in the cytosol, and spreads to neighboring cells using actin-based motility (9). The virulence factors employed by *L. monocytogenes* to successfully infect a host are all transcriptionally regulated by the master virulence regulator PrfA, which is itself redox-regulated (10). In addition to virulence factors, PrfA regulates genes that enhance *L. monocytogenes* resistance to peroxide (11), further indicating that peroxide stress is relevant during infection. We became particularly interested in the peroxide detoxification strategy employed by *L. monocytogenes* during infection when previous work revealed that the single catalase produced by the bacterium is required for aerobic growth, but dispensable during infection (12). These results led to the hypothesis that *L. monocytogenes* must produce other redundant peroxidases in order to survive the respiratory burst of the macrophage phagosome. Here, we evaluated the expression and essentiality of all peroxidase-encoding genes during *L. monocytogenes* growth *in vitro* and during infection of murine cells in tissue culture.

## RESULTS

### *L. monocytogenes* encodes 9 predicted peroxidases

*In silico* analysis of the *L. monocytogenes* proteome identified 9 proteins with predicted peroxidase activity (Table 1). Peroxidase proteins belong to two families: heme-peroxidases and non-heme peroxidases (13). Of the heme peroxidases, *L. monocytogenes* encodes a heme-dependent catalase (Kat), a coproheme decarboxylase (ChdC, formerly HemQ), and a DyP-type peroxidase (Lmo0367). The non-heme peroxidases include the peroxiredoxin family (Thiol Prx superfamily), which is characterized by highly conserved peroxidatic cysteine residues (6). In this family, *L. monocytogenes* encodes Lmo0983, AhpA (formerly AhpC or Prx), Tpx, Lmo1609, and OhrA. Finally, Fri is a non-heme bacterial ferritin classified as an oxidoreductase by the RedoxiBase database (13).

**Table 1.**
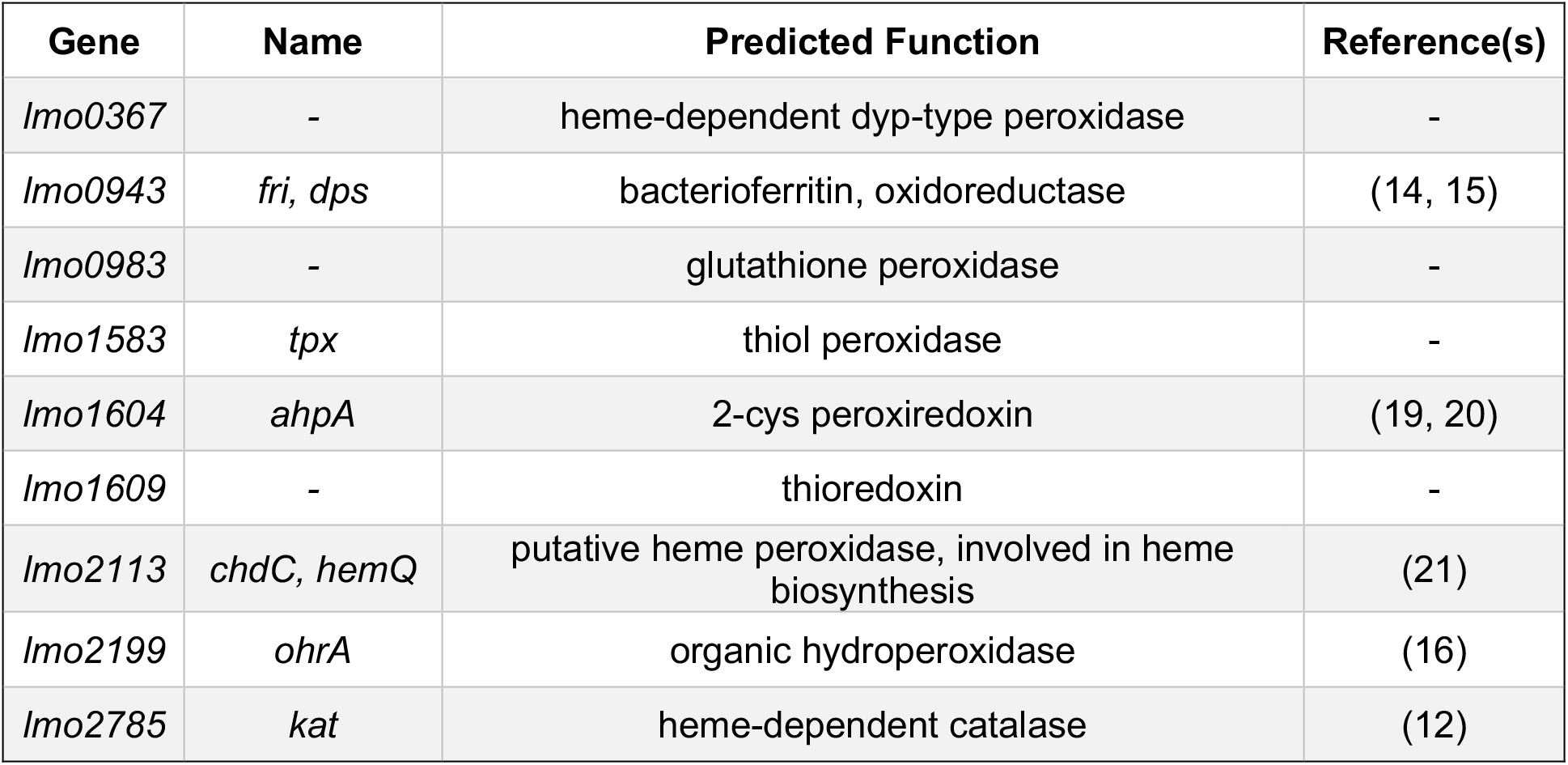
Predicted peroxidases encoded by *L. monocytogenes*

Prior research on the role of *L. monocytogenes* peroxidases is limited. The most well-studied is the ferritin protein Fri (formerly Dps), which was shown to be important for the acute peroxide stress response, long term stationary phase survival, and adaptation to shifting growth conditions (14, 15). A *L. monocytogenes* strain lacking *ohrA* is more sensitive to a variety of oxidative stressors and is attenuated during infection (16). The most dramatic phenotype associated with a *L. monocytogenes* peroxidase mutant is exhibited by strains lacking *kat*, which replicate aerobically to mid-log at the same rate as wild type (wt) but then succumb to endogenously produced peroxide toxicity (12). However, *kat* is not required for intracellular replication in macrophages or for virulence in a murine model of infection and catalase-deficient strains have been isolated from infected humans (12, 17, 18). Based on the important role of peroxide in the innate immune response and the limited research on *L. monocytogenes* peroxidases, we sought a more holistic picture of the role of peroxidases in aerobic growth and virulence.

### Expression of peroxidase-encoding genes

To investigate which peroxidase enzymes may be important during *L. monocytogenes* aerobic growth, we first evaluated gene expression in rich media by quantitative reverse transcriptase PCR (qPCR). In these experiments, bacteria were grown in tryptic soy broth (TSB) which lacks glutathione and heme, both of which influence the redox environment of bacterial cultures. Expression of each putative peroxidase-encoding gene was evaluated in wt and Δ*kat* strains and expression of transcripts was normalized to wt at 2 hours post-inoculation into shaking flasks. The Δ*kat* mutant replicated to mid-log, but stopped growing and began to die 8 hours post-inoculation (Fig. 1A and reference 12). This requirement for Kat activity in early stationary phase corresponds to the peak expression of *kat* in wt *L. monocytogenes* (Fig. 1B). Accordingly, we found that gene expression could not be reliably measured in Δ*kat* 8 hours post-inoculation due to bacterial death. Analyzing transcript abundance of the other peroxidase-encoding genes revealed that lmo0367, lmo0983, ahpA, and lmo1609 did not increase appreciably over time in the wt or Δ*kat* background strains (Fig. 1C). In contrast, tpx expression was elevated early in *L. monocytogenes* lacking *kat* compared to wt, although this was not statistically significant (Fig. 1D, *p* = 0.2). The gene encoding Fri was increased in wt *L. monocytogenes* 8 hours post-inoculation, similarly to *kat* expression. Expression of *chdC* exhibited a small, but statistically significant increase initially in *L. monocytogenes* lacking *kat* compared to wt and was significantly increased in wt after 6 hours of aerobic growth (Fig. 1D). The gene encoding OhrA exhibited maximal expression 2 hours post-inoculation and decreased dramatically over time in both strains. Taken together, these data demonstrated that *kat* and *fri* share similar expression dynamics during aerobic growth.

**Figure 1.**
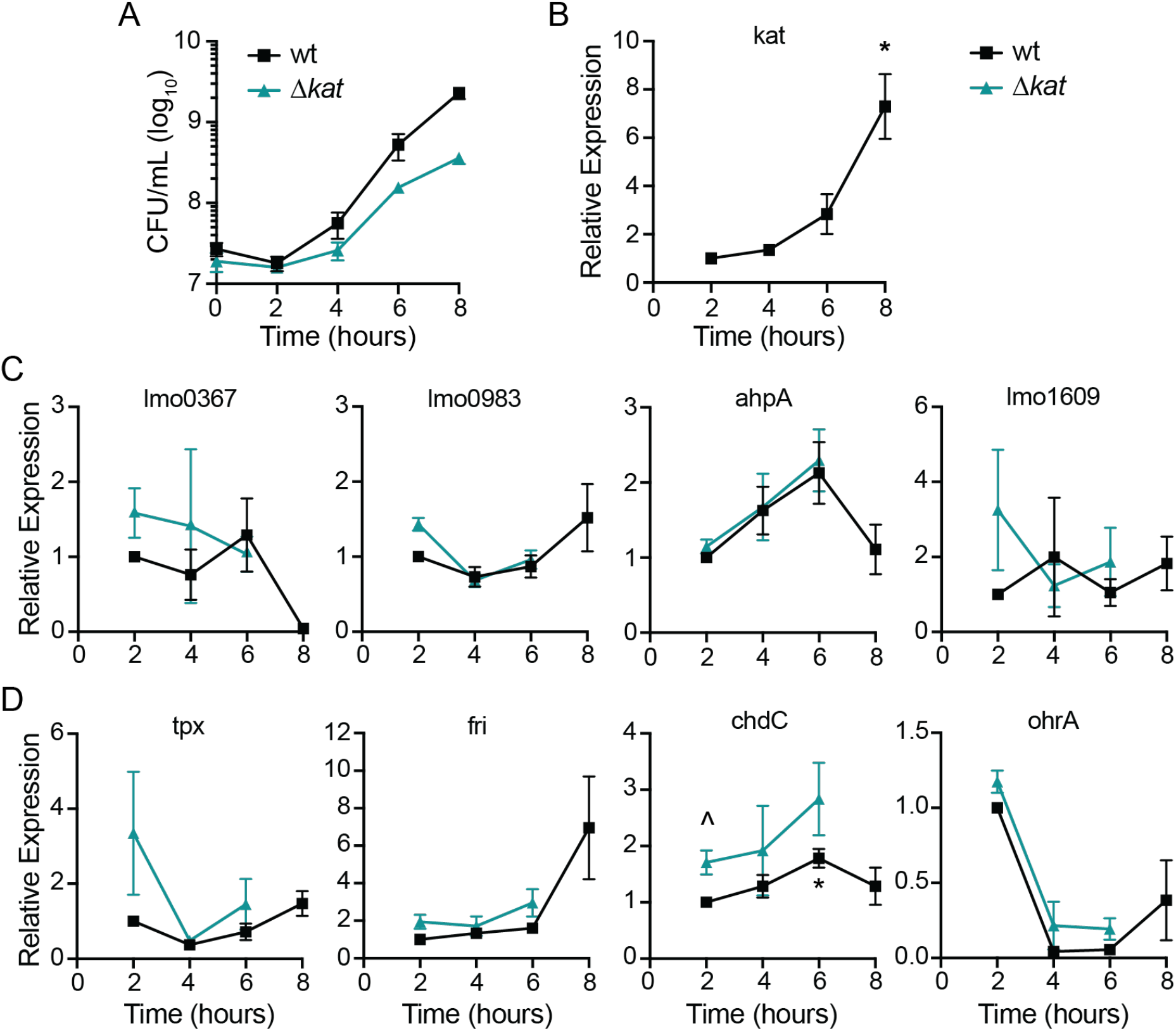
Expression of genes encoding putative peroxidases during aerobic growth. **A.** Aerobic growth of wt and Δ*kat* in shaking flasks was measured by plating for CFU and incubating the plates anaerobically. Data are the means and standard error of the means (SEM) for three biological replicates. **B - D.** Relative expression of putative peroxidase-encoding genes over time in both wt and Δ*kat*, as grown in *A*. Expression was normalized to wt expression at 2 hours. Data are the means and standard error of the means (SEMs) of at least three biological replicates. *p* values were calculated using a heteroscedastic Student’s *t* test. * *p* < 0.05 expression compared to wt at 2 hours; ^ *p* < 0.05 expression in Δ*kat* compared to wt at that time point.

We next measured expression of these peroxidase-encoding genes during infection using fluorescent transcriptional reporters. Strains were engineered to express *rfp* from the native promoter of each peroxidase-encoding gene and *gfp* was expressed constitutively. J774 macrophages were infected for 1 hour before gentamicin was added to the media to eliminate extracellular bacterial growth. Flow cytometry was performed 6 hours post-infection and infected cells were identified by GFP fluorescence compared to an uninfected control. Cells infected with the reporter strains for expression of *fri*, *ahpA*, and *kat* exhibited significantly increased RFP production compared to the background (Fig. 2). These results suggested that only *fri, ahpA*, and *kat* are significantly expressed during macrophage infection and therefore might have a role in virulence.

**Figure 2.**
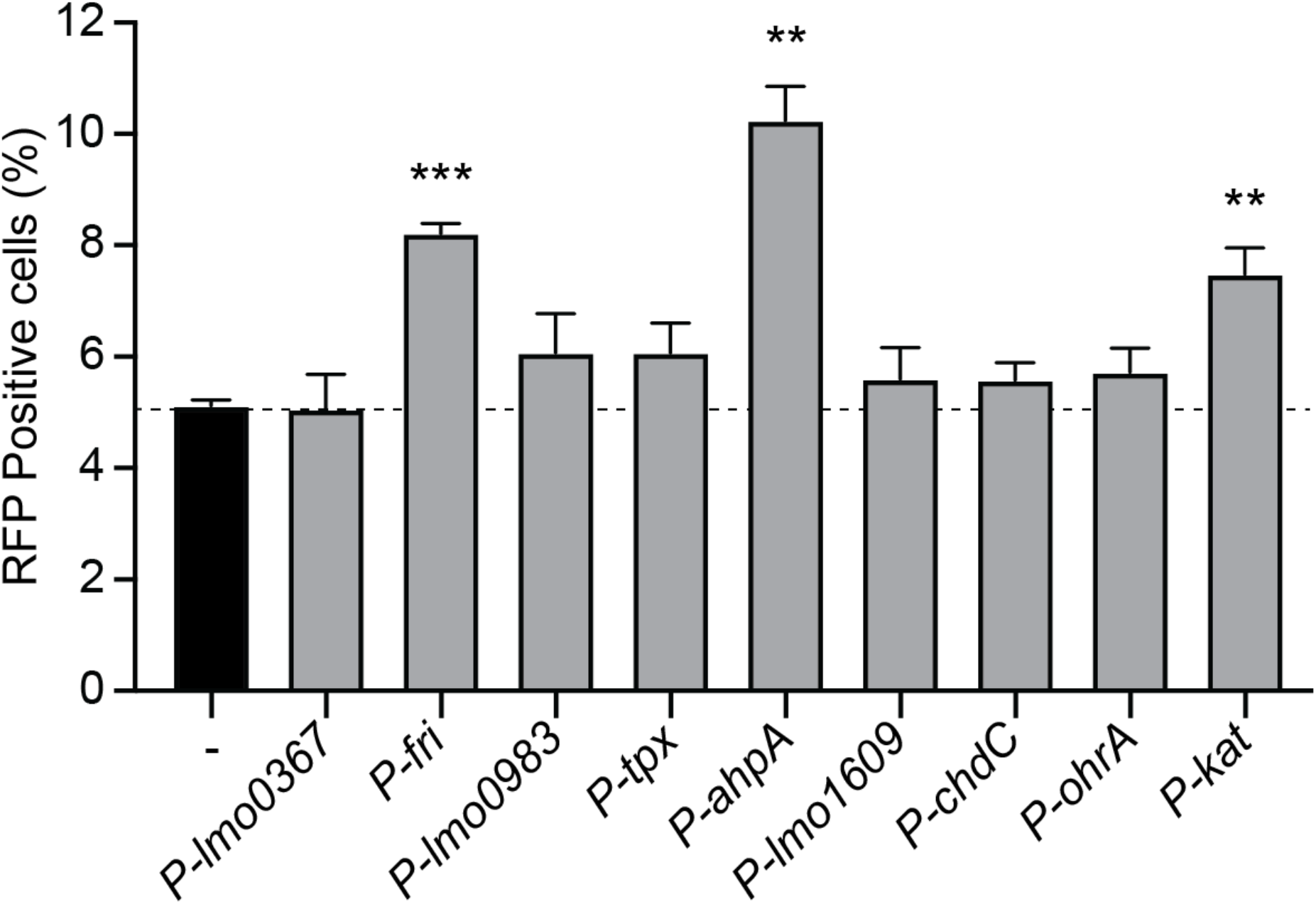
Intracellular expression of peroxidase-encoding genes. J774 macrophages were infected with each reporter strain, which expressed *rfp* from the indicated promoter and constitutive *gfp*. Cells were infected for six hours and then analyzed by flow cytometry. The dotted line indicates background RFP fluorescence. Data are the means and SEMs of three biological replicates. *p* values were calculated using a heteroscedastic Student’s *t* test comparing each mutant to wt. ** *p* < 0.01; *** *p* < 0.001.

### Growth of peroxidase mutants in broth

Previous work demonstrated that *L. monocytogenes* requires *kat* to detoxify endogenously-produced peroxide during aerobic growth (12). Here, we sought to test if additional peroxidases are important for aerobic growth. For this analysis, each potential peroxidase-encoding gene was deleted by allelic exchange while growing anaerobically to prevent oxygen-mediated toxicity. Based on the redundant roles of Kat and Ahp in other bacteria (7, 22), we also generated a Δ*kat*Δ*ahpA* double mutant. Anaerobic bacterial overnight cultures were diluted into rich media and grown aerobically in shaking flasks. Serial dilutions were performed to enumerate CFU over time and the plates were incubated anaerobically to promote growth of potentially oxygen-sensitive strains.

Growth analyses revealed that several peroxidase-encoding genes were dispensable for aerobic replication, including: *lmo0367, lmo0983, tpx, ohrA, lmo1609*, and *ahpA* (Fig. 3A and 3B). In contrast, *kat* was required for aerobic growth and began to die upon entry into stationary phase (Fig. 3B), as previously reported (12). The Δ*kat*Δ*ahpA* double mutant was even more sensitive to oxygen, as we observed a significant attenuation in survival at 7 and 9 hours post-inoculation compared to the single mutant lacking *kat* (Fig. 3B). These data indicate that AhpA is functional and important for aerobic replication in the absence of catalase.

**Figure 3.**
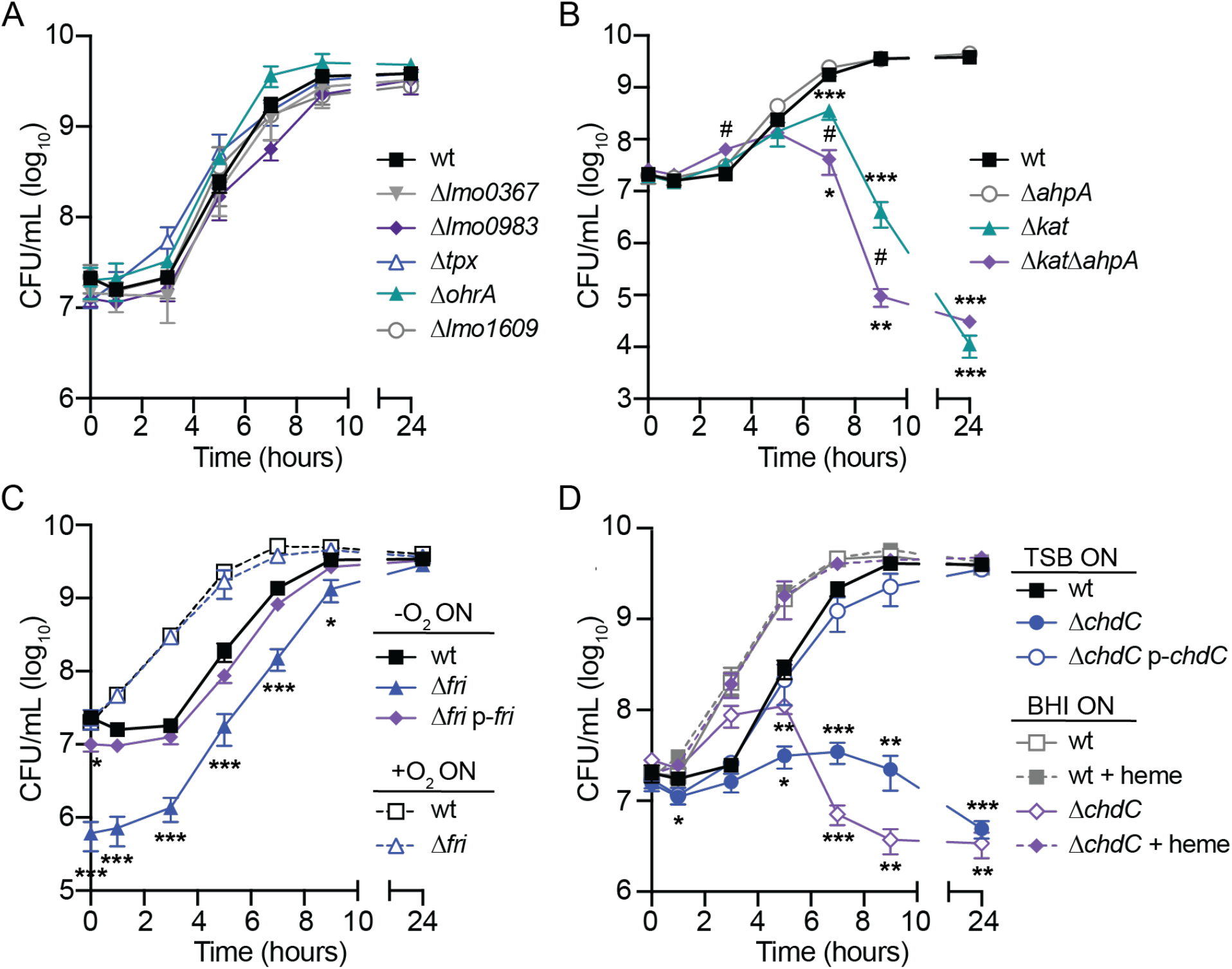
Aerobic growth of peroxidase mutants. **A.** Growth curves of strains that replicate at the same rate as wt (*p* > 0.05 at each time point). **B.** Growth curves comparing Δ*ahpA*, Δ*kat*, and Δ*kat*Δ*ahpA*. # indicates significant difference between Δ*kat* and Δ*kat*Δ*ahpA* (*p* < 0.05). **C.** Growth curves of strains grown anaerobically (solid lines) or aerobically (dotted lines) overnight before back-diluting into shaking flasks. **D.** Strains were grown anaerobically overnight in TSB or BHI and then diluted into TSB with or without exogenous heme (5 μM). In all panels, data are the mean and SEMs of three biological replicates. *p* values were calculated using a heteroscedastic Student’s *t* test comparing each mutant to wt grown in the same conditions. * *p* < 0.05; ** *p* < 0.01; *** *p* < 0.001.

Interestingly, we observed a dramatic phenotype for the *L. monocytogenes* strain lacking *fri* (Fig. 3C). Despite normalizing the anaerobic overnight cultures by optical density, we consistently observed a 1- to 2-log defect in Δ*fri* CFU at the initial time point and this defect continued throughout growth until stationary phase (Fig. 2C). This aerobic growth defect could be complemented by providing a copy of *fri in trans* using the integrative plasmid pPL2 (p-*fri*) (23). Previous work suggested that *fri* is important for *L. monocytogenes* adaptation to stress conditions, including nutritional stress and temperature shifts (15). We therefore hypothesized that Δ*fri* was unable to rapidly adapt from hypoxia to aerobic growth. In support of this hypothesis, we found that Δ*fri* incubated aerobically overnight grew similarly to wt after diluting into shaking flasks (Fig. 3C, dotted lines). Consequently, the Δ*fri* mutant was incubated overnight aerobically for all subsequent experiments.

Finally, we observed the Δ*chdC* mutant did not replicate when diluted into shaking flasks and this growth defect was genetically complemented by expressing *chdC in trans* (p-*chdC*, Fig. 3D). This finding is in agreement with published work reporting that *chdC* is essential in *L. monocytogenes* (24). ChdC catalyzes the final step of heme biosynthesis and therefore the Δ*chdC* mutant lacks both peroxidase activity and endogenous heme (21). To distinguish which function is important for aerobic growth, we attempted to chemically complement Δ*chdC* growth by supplementing with exogenous heme. However, addition of exogenous heme is toxic unless bacterial cultures are pre-exposed to low levels of heme (25). Therefore, bacteria were first grown overnight in brain heart infusion (BHI) which contains heme and then diluted into TSB containing or lacking heme (5 μM). Exogenous heme fully restored growth of Δ*chdC* to that of wt (Fig. 3D, dotted lines), suggesting that the aerobic growth defect of *L. monocytogenes* lacking *chdC* is primarily due to a lack of heme.

### Peroxidases important for acute peroxide toxicity

We next investigated which peroxidases are required for detoxifying acute peroxide stress. Bacteria were grown overnight aerobically, as *L. monocytogenes* does not produce detectable peroxidase activity after anaerobic culture (12). Accordingly, only the mutants which replicated aerobically could be tested in this experiment. Each strain was grown to early logarithmic phase (OD_600_ = 0.6) before hydrogen peroxide (120 mM) was added for 1 hour. At this concentration of peroxide, wt decreased 1.7-fold over one hour and most mutants exhibited similar resistance (Fig. 4). In contrast, both Δ*fri* and Δ*ahpA* were rapidly killed and no bacteria were detected at 30 or 60 minutes. These results are consistent with published reports demonstrating *fri* and *ahpA* are important for survival in the presence of peroxide (14, 19, 20), and reveal that none of the other peroxidases tested provide non-redundant protection from peroxide under these conditions.

**Figure 4.**
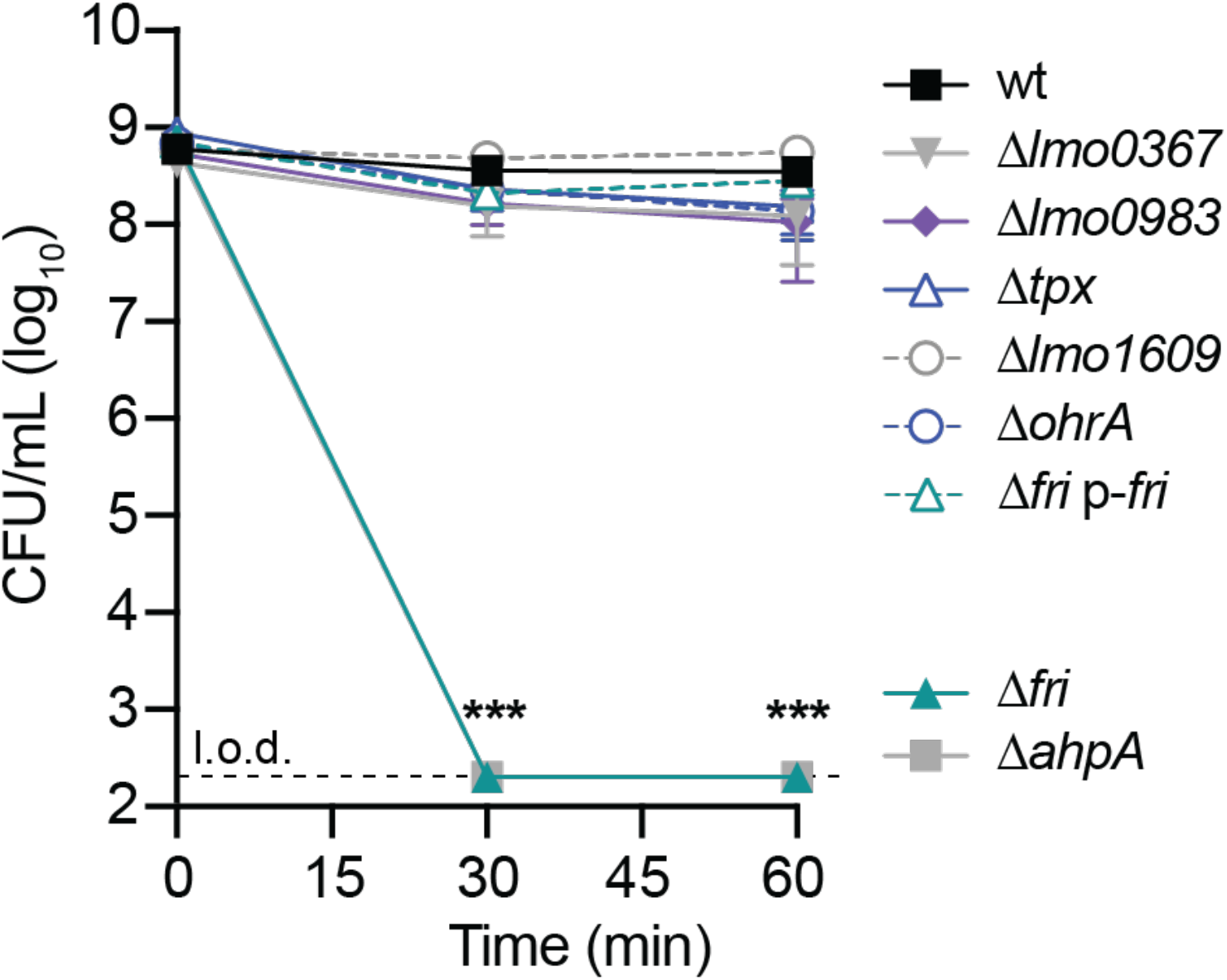
Acute peroxide toxicity. Bacteria were grown aerobically to mid-log before hydrogen peroxide (120 mM) was added. The dotted line indicates the limit of detection (l.o.d.). Data are the means and SEMs of four biological replicates. *p* values were calculated using a heteroscedastic Student’s *t* test comparing each mutant to wt. *** *p* < 0.001.

### Intracellular growth of peroxidase mutants

During infection of macrophages, *L. monocytogenes* resides briefly in the phagosome where it is bombarded by hydrogen peroxide produced by the host respiratory burst (1). Previous work demonstrated that Kat-mediated peroxide detoxification is not required for intracellular survival or growth in macrophages (12). To test which peroxidases are important intracellularly, we infected bone marrow-derived macrophages (BMDMs) with each strain at an MOI of 0.1 and enumerated CFU over time by plating anaerobically. All the mutants replicated similarly to wt in resting BMDMs (data not shown). To ensure a robust innate immune response, we next infected BMDMs that were pre-treated with interferon gamma (IFNγ) to activate the host respiratory burst (26). In these experiments, the majority of peroxidase mutants grew similarly to wt (Fig. 5A). Surprisingly, the Δ*kat*Δ*ahpA* double mutant exhibited a significant 5-8-fold increase in CFU at each time point compared to wt (Fig. 5B). In contrast, the Δ*fri* strain was significantly attenuated for intracellular replication in activated BMDMs and this defect was genetically complemented by providing *fri in trans* (Fig. 5B).

**Figure 5.**
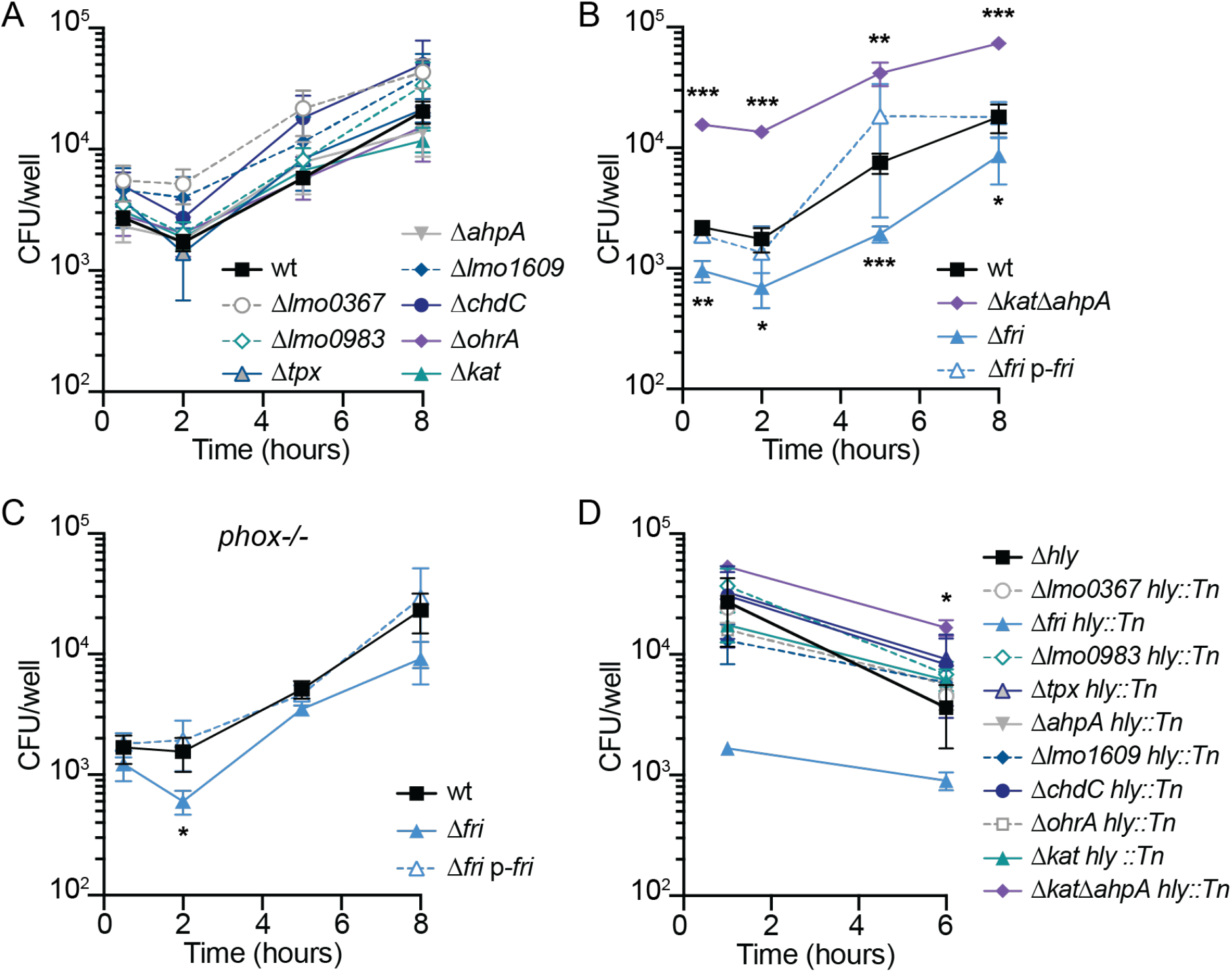
Intracellular survival and growth of peroxidase mutants in activated BMDMs. **A.** Intracellular growth curves of mutant strains that replicated at the same rate as wt (*p* > 0.05 at each time point). **B. C.** Intracellular growth curves in *phox*^-/-^ BMDMs which lack NADPH oxidase. **D.** Survival of *hly* mutants trapped in the vacuoles of activated BMDMs. Although the error bars are too small to be visible, the Δ*fri hly::Tn* strain is not significantly different from wt at 1 hour (*p* = 0.06). In all panels, data are the mean and SEMs of at least three independent experiments. *p* values were calculated using a heteroscedastic Student’s *t* test comparing each mutant to wt grown in the same conditions. * *p* < 0.05; ** *p* < 0.01; *** *p* < 0.001.

Due to the role of Fri in iron storage, we sought to assess whether the growth defect of Δ*fri* was due to a lack of iron scavenging or peroxidase protection. To that end, intracellular growth curves were performed in gp91^*phox*-/-^ (*phox*^-/-^) macrophages which lack the NADPH oxidase responsible for the host oxidative bursts and thus mimics the most common genetic defect observed in humans with CGD (3). In these immunodeficient cells, the Δ*fri* mutant exhibited a significant defect 2 hours post-infection, but the intracellular bacterial burden was similar to that of wt and the complemented strain at every other time point (Fig. 5C). Taken together, these results demonstrate that while the Δ*kat*Δ*ahpA* mutant has an unexpected advantage during infection, *fri* is required for efficient infection of activated macrophages and this is at least partially alleviated in the absence of the host respiratory burst.

One hypothesis to explain the dispensability of the majority of peroxidases for intracellular growth (Fig 5A), is that the host respiratory burst is ineffective against *L. monocytogenes* because the secreted pore-forming toxin LLO allows for rapid escape from the vacuole. To examine which peroxidases may be important specifically in the vacuolar environment, the gene encoding LLO (*hly*) was disrupted in each mutant background and survival in the vacuole of IFNγ-activated BMDMs was evaluated over time. In agreement with the intracellular growth curves, the majority of peroxidase mutants survived in the vacuole for 6 hours at rates similar to wt (Fig. 5D). The exceptions were the Δ*kat*Δ*ahpA* mutant, which survived significantly better than wt, and Δ*fri*, which exhibited a ~1-log decrease in CFU at the earliest time point, although this was not statistically significant (*p* = 0.06). Together, these data demonstrate that the only peroxidase important for vacuolar survival and intracellular growth in activated BMDMs is *fri*.

### Intercellular spread

We next sought to test the role of each peroxidase in virulence using a plaque assay, which is a measure of intracellular growth and cell-to-cell spread that is highly correlated with virulence in a murine model of infection (16, 27). In this assay, a monolayer of L2 murine fibroblasts is infected and immobilized in agarose containing gentamicin to kill extracellular bacteria. Three days post-infection the living cells are stained with neutral red and the area of the plaques formed by *L. monocytogenes* are analyzed as a measure of intercellular spread. In this assay, all of the mutant strains formed plaques similar in size or larger than those formed by wt (Fig. 6A). Although the Δ*kat*Δ*ahpA* double mutant formed plaques of a similar size as wt, the mutant exhibited a dramatic advantage at invading host cells. The Δ*kat*Δ*ahpA* formed approximately 5 times more numerous plaques than wt (Fig. 6B). This is consistent with the increased bacterial burden observed in IFNγ-activated BMDMs at the earliest time points (Fig. 5B), and suggests that Δ*kat*Δ*ahpA* is better able to invade host cells.

**Figure 6.**
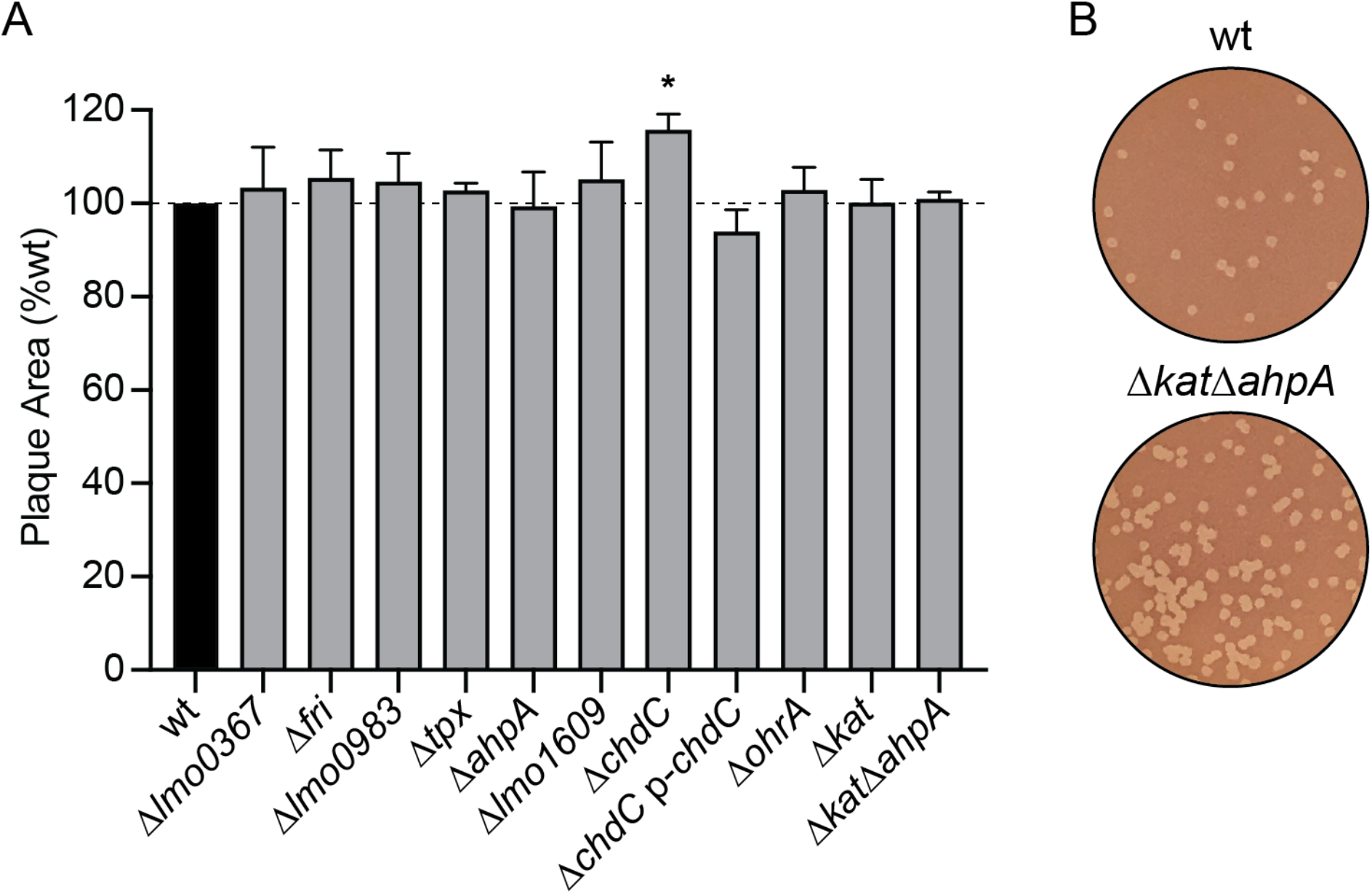
Intercellular spread of peroxidase mutants. **A.** Plaque formation in L2 fibroblasts was evaluated for each strain. Data are graphed as mean and SEMs. The dotted line signifies the 100% of wt. **B.** Representative images of plaques demonstrating the greater number of plaques formed by the Δ*kat*Δ*ahpA* mutant compared to wt.

## DISCUSSION

Bacterial pathogens must detoxify endogenously produced ROS and exogenous sources of oxidative stress during infection. Hydrogen peroxide is particularly dangerous, as it is an uncharged molecule that can penetrate membranes (5). In this study, we performed the first comparison of the expression and essentiality of all proteins with predicted peroxidase activity in *L. monocytogenes*. Our results revealed that *kat* and *chdC* are required for aerobic growth in rich media, while *ahpA* and *fri* are required to detoxify acute peroxide stress *in vitro*. The strain lacking both *kat* and *ahpA* exhibited the most severe aerobic growth defect, but had a surprising advantage invading and surviving in host cells. While *fri, ahpA*, and *kat* were expressed during intracellular growth in macrophages, only *fri* was required for survival in INFγ-activated BMDMs. Considering the host respiratory burst assaults invading bacteria with up to 100 μM peroxide in the phagosome (7), our results suggest a high degree of redundancy in the *L. monocytogenes* peroxide stress response.

Transcript analysis during aerobic growth revealed that *L. monocytogenes* expression of peroxidase-encoding genes was not dramatically altered in the *kat*-deficient strain compared to wt. This was unexpected, as *B. subtilis* strains lacking *katA* or *ahpC* experience peroxide stress during aerobic growth that induces the PerR (peroxide stress response) regulon (28, 29). Similarly, the *E. coli* OxyR regulon is activated by the peroxide that accumulates when Hpx-cells (Δ*katE*Δ*katG*Δ*ahpCF*) are grown aerobically (7, 30). Thus, one interpretation of our results is that *L. monocytogenes* Δ*kat* succumbs to peroxide-mediated toxicity before the PerR regulon can be effectively activated (8). However, the Δ*kat*Δ*ahpA* double mutant is even more susceptible to endogenous redox stress *in vitro*, but undergoes undefined regulatory changes that result in increased invasion, survival, and replication in activated macrophages. Ongoing research is aimed at deciphering the regulatory changes that occur in the absence of *kat* and *ahpA* that lead to increased virulence.

Lmo1604 is annotated as an alkyl hydroperoxide reductase (Ahp) based on similarity to other enzymes. The classical Ahp system is comprised of two components: the peroxiredoxin (AhpC) and a dedicated reductase (AhpF) that reduces and recycles AhpC and is typically encoded by a sequence adjacent to *ahpC* (31). Many organisms encode multiple Ahp proteins, however *L. monocytogenes* encodes only Lmo1604 and there is no adjacent reductase. *L. monocytogenes* Lmo1604 shares 75% and 33% identity with *B. subtilis* AhpA and AhpC, respectively. While two previous publications referred to this protein as Prx (19, 20), herein we refer to Lmo1604 as AhpA to reflect its similarity to *B. subtilis* AhpA and the fact that it is a noncanonical Ahp that lacks a dedicated reductase. We found that *ahpA* is expressed during intracellular replication, but is not required for growth in IFNγ-activated macrophages or prolonged survival in the vacuole. These results are consistent with published work showing the Δ*ahpA* mutant had no defect in macrophages or during infection of mice (20). However, *ahpA* was required for surviving acute peroxide stress *in vitro* and was important for detoxifying endogenous peroxide during aerobic replication in the absence of catalase. Together, these results suggest a role for *L. monocytogenes* AhpA in peroxide detoxification that is masked in wt cells by the activity of catalase. Interestingly, the opposite is true in *E. coli* where Ahp is the primary scavenger of endogenous peroxide and catalase is important only in the presence of high concentrations of peroxide (30).

Hydrogen peroxide toxicity and iron are inextricably linked due to the Fenton reaction, in which ferrous iron reacts with hydrogen peroxide to generate hydroxyl radicals that damage DNA (7). Unsurprisingly, several of the peroxidases in this study are also involved in maintaining iron homeostasis in the cell, including: the heme-dependent peroxidase Lmo0367, the bacterial ferritin Fri, the terminal heme biosynthesis enzyme ChdC, and the heme-dependent catalase Kat. Additional studies are necessary to assess the roles of these enzymes in iron influx, oxidation, and storage in response to oxidative stress.

One method used by bacteria to mitigate the danger of free iron and the Fenton reaction is to sequester iron within bacterial ferritin proteins, which are multimeric protein shells that can store up to 4500 iron atoms. The amino acid sequence of *L. monocytogenes* Fri is identical to that of the *L. innocua* Dps protein, which has been biochemically characterized as a bacterial ferritin. *L. innocua* Dps was named due to its structural similarity to Dps family proteins (<u>D</u>NA-binding <u>p</u>rotein from <u>s</u>tarved cells), although it does not bind DNA (32). *L. innocua* Dps has a smaller internal diameter than typical ferritins and therefore can store only ~400 iron atoms per shell. The protection afforded by *L. innocua* Dps is due to the hydrogen peroxide-mediated iron oxidation, which occurs rapidly and in a manner that prevents Fenton chemistry (32, 33). Based on the identity of the proteins, we predict *L. monocytogenes* Fri has a similar function and uses peroxide to oxidize and store iron. It is therefore not possible to distinguish between the importance of the peroxidase activity of Fri and its role in iron homeostasis in the cell.

The role of Fri in *L. monocytogenes* has been investigated previously by several groups and in fact, the protein encoded by *lmo0943* has been given multiple different names, including: Flp (<u>f</u>erritin-<u>l</u>ike <u>p</u>rotein) (34), Frm (<u>f</u>e<u>r</u>ritin-like protein from *L. <u>m</u>onocytogenes*) (35), and Frl (<u>f</u>e<u>r</u>ritin-like protein in *<u>L</u>isteria* species) (36). Here, we kept the more common name Fri (14, 15, 37). Our results are consistent with the literature showing that *fri* expression increases 8-fold upon entry into stationary phase and a Δ*fri* mutant is more sensitive to peroxide stress when treated in log phase (14, 37). In addition, we identified a critical role for Fri in the transition from anaerobiosis to aerobic replication. We also determined that *fri* expression is increased during intracellular growth and accordingly, *fri* is required for growth in activated BMDMs. This requirement was partially alleviated in host cells incapable of mounting an effective respiratory burst, suggesting that host-derived peroxide contributes to limiting Δ*fri* intracellular replication.

*L. monocytogenes* and *Staphylococcus aureus* mutants deficient in heme biosynthesis enzymes form small colonies on solid media and grow poorly in aerobic liquid cultures (12, 38). While *chdC* was previously reported to be essential (24), we generated a Δ*chdC* mutant in anaerobic growth conditions and observed that it was indeed unable to replicate aerobically in the absence of exogenous heme. This severe growth defect resembled that of *L. monocytogenes* Δ*hemEH*, which also cannot produce heme (12). As our data attribute the Δ*chdC* growth defect to the heme deficiency, it is unlikely that ChdC peroxidase activity plays a primary role in endogenous peroxide detoxification. However, our results raise an interesting question: why is heme biosynthesis required for *L. monocytogenes* aerobic growth? Heme is an essential cofactor for cytochrome oxidases of the electron transport chain and therefore, *S. aureus hem* mutants are impaired for growth due to a lack of aerobic respiration (38, 39). However, *L. monocytogenes* lacking one or both terminal cytochrome oxidases are only moderately impaired for aerobic growth as compared to the complete lack of replication observed for Δ*chdC* and Δ*hemEH* mutants (12, 40). Interestingly, *S. aureus* requires heme biosynthesis for virulence in murine models of acute infection (41). In contrast, *L. monocytogenes* lacking *hemEH* or *chdC* are not defective for intracellular replication and in fact, exhibit increased intercellular spread compared to wt (reference 12 and N.H.S. unpublished observations). These results suggest that either heme biosynthesis is dispensable for *L. monocytogenes* pathogenesis or that exogenous host-derived heme can support growth of Δ*chdC* during infection. Ongoing research aims to determine the role of heme in *L. monocytogenes* aerobic growth and virulence.

In our assays, we did not observe phenotypes for *L. monocytogenes* lacking *lmo0367, lmo0983, tpx, lmo1609*, or *ohrA* and little is known about their functions. Lmo0367 shares 52% amino acid identity with the *B. subtilis* YwbN/EfeB protein, a DyP-type peroxidase that is transported as a folded protein across the cytoplasmic membrane via the twin-arginine translocation (Tat) pathway (42, 43). While protein localization and function of *L. monocytogenes* Lmo0367 have not been examined, the corresponding gene was found to be regulated by the ferric uptake regulator Fur and expression was consequently decreased in response to heme stress (44, 45). A screen for *L. monocytogenes* genes important for osmotic stress and desiccation identified a transposon in *lmo0983*, encoding a glutathione peroxidase, although this mutant was not further characterized (46). Tpx and Lmo1609 share 63% and 58% amino acid identity with their respective homologues in *B. subtilis* and both are activated by Spx in that organism, supporting their role in the oxidative stress response (47). The *L. monocytogenes ohrA* mutant was previously found to be attenuated in a plaque assay and for intracellular replication in BMDMs (16). In this study, the Δ*ohrA* strain was not attenuated and we hypothesize this discrepancy is due to the fact that herein, the bacteria were grown overnight anaerobically before infecting cells, whereas previous experiments grew bacteria aerobically. Future research will decipher the roles of these understudied peroxidases in *L. monocytogenes* and other Firmicutes.

Oxidative stress is abundant in the environment during aerobic growth and during infection. In this work we focused on peroxide stress, although superoxide is also generated endogenously and encountered exogenously in host phagocytes (1). *L. monocytogenes* produces a single manganese-dependent superoxide dismutase (MnSOD) that is required for infection (48, 49). MnSOD converts the superoxide produced by NADPH oxidase to hydrogen peroxide, which then needs to be detoxified by catalases and peroxidases. Our results demonstrate that the majority of peroxidases are individually dispensable for infection of mammalian cells and suggest redundancy in these antioxidants. Identifying expression changes in the Δ*kat*Δ*ahpA* strain will reveal the compensatory factors allowing this double mutant to more efficiently infect macrophages, despite the *in vitro* sensitivity of this strain to peroxide. Future investigations will build on the results described herein to provide further insight into the *L. monocytogenes* peroxide detoxification strategy during infection.

## MATERIALS AND METHODS

### Ethics Statement

This study was carried out in strict accordance with the recommendations in the Guide for the Care and Use of Laboratory Animals of the National Institutes of Health. All protocols were reviewed and approved by the Institutional Animal Care and Use Committee at the University of Washington (protocol 4410-01).

### Bacterial strains and culture conditions

*L. monocytogenes* strains were derived from the wt strain 10403S and are listed in Table 2. *E. coli* strains are listed in Table 3. *L. monocytogenes* was cultured in either tryptic soy broth (TSB) or brain heart infusion (BHI), aerobically in the dark shaking at 37°C unless other conditions were noted. Anaerobic conditions were established by growing bacteria in closed containers with GasPak EZ Anaerobe Gas Generating Pouches (Becton Dickinson) or placing cultures in degassed media inside of a closed-system anaerobic chamber (Don Whitley Scientific A35 Anaerobic Workstation). Unless otherwise stated, the chemicals used were purchased from Sigma Aldrich. The following concentrations of antibiotics were used: streptomycin, 200 μg mL^-1^; chloramphenicol, 10 μg mL^-1^ (*E. coli*), 7.5 μg mL^-1^ (*L. monocytogenes*); carbenicillin, 100 μg mL^-1^; erythromycin, 1μg mL^-1^; and tetracycline, 1 μg mL^-1^.

**Table 2.** *L. monocytogenes* strains used in this study.

| Strain | Description | Reference |
| --- | --- | --- |
| MLR-L001 | 10403S | (51, 52) |
| MLR-L948 | $\Delta lmo0367$ | This work |
| MLR-L960 | $\Delta fri$ (*lmo0943*) | This work |
| MLR-L954 | $\Delta lmo0983$ | This work |
| MLR-L959 | $\Delta tpx$ (*lmo1583*) | This work |
| MLR-L879 | $\Delta ahpA$ (*lmo1604*) | This work |
| MLR-L953 | $\Delta lmo1609$ | This work |
| MLR-L944 | $\Delta chdC$ (*lmo2113*) | This work |
| MLR-L174 | $\Delta ohrA$ (*lmo2199*) | (16) |
| MLR-L828 | $\Delta kat$ (*lmo2785*) | (12) |
| MLR-L880 | $\Delta kat\Delta ahpA$ | This work |
| MLR-L971 | $\Delta fri$ p-*fri* | This work |
| MLR-L850 | pPL1.pHyper-*gfp* (pH-*gfp*) | This work |
| MLR-L984 | pH-*gfp* pHyper-*rfp* | This work |
| MLR-L985 | pH-*gfp* P-*lmo0367-rfp* | This work |
| MLR-L986 | pH-*gfp* P-*fri-rfp* | This work |
| MLR-L987 | pH-*gfp* P-*lmo0983-rfp* | This work |
| MLR-L988 | pH-*gfp* P-*tpx-rfp* | This work |
| MLR-L983 | pH-*gfp* P-*ahpA-rfp* | This work |
| MLR-L989 | pH-*gfp* P-*lmo1609-rfp* | This work |
| MLR-L982 | pH-*gfp* P-*chdC-rfp* | This work |
| MLR-L990 | pH-*gfp* P-*ohrA-rfp* | This work |
| MLR-L981 | pH-*gfp* P-*kat-rfp* | This work |
| MLR-L991 | *hly::himar1* (Tn) | (10) |
| MLR-L992 | $\Delta lmo0367$ *hly::Tn* | This work |
| MLR-L1000 | $\Delta fri$ *hly::Tn* | This work |
| MLR-L995 | $\Delta lmo0983$ *hly::Tn* | This work |
| MLR-L996 | $\Delta tpx$ *hly::Tn* | This work |
| MLR-L999 | $\Delta ahpA$ *hly::Tn* | This work |
| MLR-L997 | $\Delta lmo1609$ *hly::Tn* | This work |
| MLR-L994 | $\Delta chdC$ *hly::Tn* | This work |
| MLR-L998 | $\Delta ohrA$ *hly::Tn* | This work |
| MLR-L993 | $\Delta kat$ *hly::Tn* | This work |
| MLR-L1001 | $\Delta kat\Delta ahpA$ *hly::Tn* | This work |

**Table 3.**
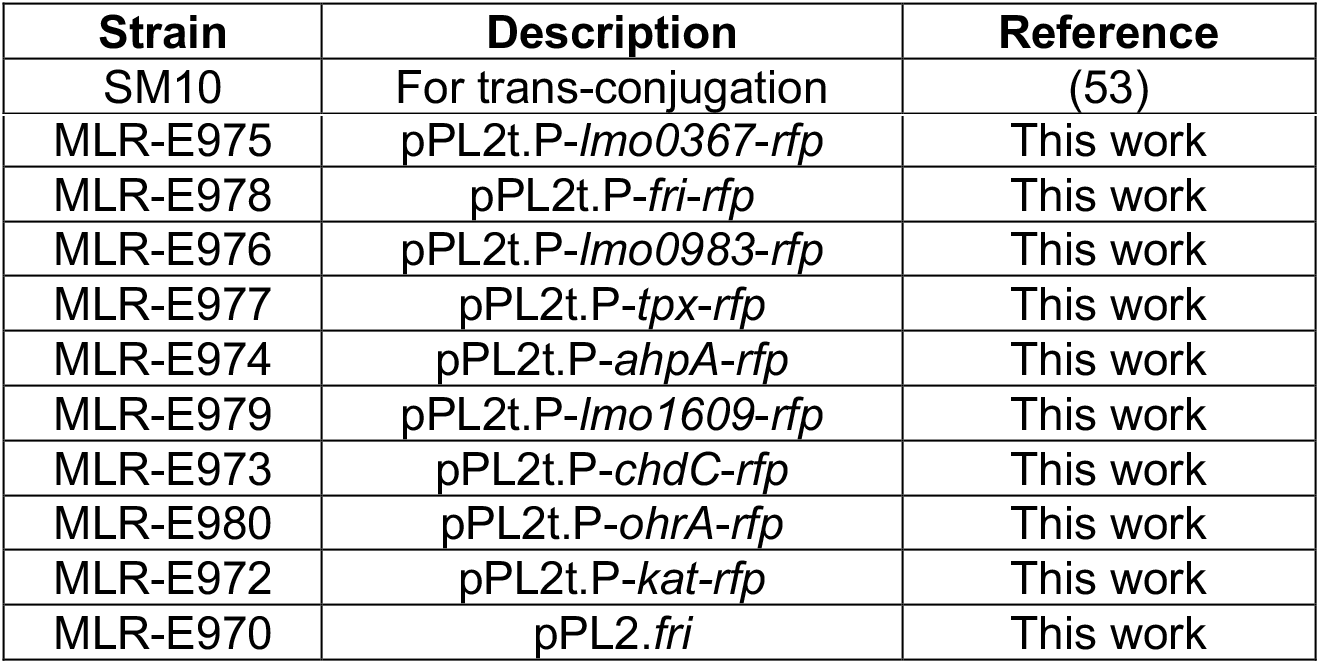
*E. coli* SM10 strains used in this study.

Tissue culture cells were routinely cultured in high-glucose Dulbecco modified Eagle medium (DMEM) at 37°C with 5.5% CO_2_. L2 fibroblasts were generated previously from L929 cells (16, 27). L2s and J774 macrophages were maintained in media containing 10% FBS (HyClone), 2 mM L-glutamine (Gibco), and 1 mM sodium pyruvate (Gibco). Bone marrow-derived macrophages (BMDMs) were derived as previously described (16, 50). Bone marrow was from C57BL/6 mice purchased from The Jackson Laboratory or from B6.129S-*Cybb^tm1Din^* (also known as gp91^*phox*-/-^) mice, a generous gift from the Fang Laboratory (University of Washington) and originally from The Jackson Laboratory. BMDMs were cultured in 20% FBS, 2 mM L-glutamine, 1 mM sodium pyruvate, BME (55 μM) and 10% M-CSF (conditioned media from 3T3 cells expressing M-CSF).

### Cloning and plasmid construction

In-frame deletions were carried out via the conjugatable suicide vector pLIM1 and allelic exchange (plasmid provided as a generous gift from Arne Rietsch, Case Western Reserve University). Complementation of in-frame deletions at ectopic loci were accomplished using pPL2 integration plasmids as previously described (23). Whole genes accompanied by promoter regions were amplified and ligated into pPL2. Constructs were transformed into *E. coli* SM10 cells and introduced into *L. monocytogenes* mutants via trans-conjugation. Complemented mutants were confirmed by antibiotic resistance and Sanger sequencing.

Fluorescent transcriptional reporters were engineered to express *gfp* constitutively from the HyPer promoter using the integrative plasmid pPL1 (16, 23). This was integrated into the chromosome of a phage-cured wt *L. monocytogenes* strain DP-L4056 (23). Next, the promoter regions of peroxidase genes were amplified and ligated to mTAG-RFP via NEBuilder HiFi DNA assembly. The mTAG-RFP was amplified from DP-L6508 (16). The promoter-*rfp* fusions were ligated into pPL2t and confirmed via PCR and Sanger sequencing. *E. coli* SM10 harboring the pPL2t.Promoter-*rfp* constructs were mated with MLR-L850 (pPL1.p*Hyper-gfp*) to generate the two-color transcriptional reporter strains. Integration was confirmed by antibiotic resistance and PCR.

Peroxidase mutants unable to escape from the vacuole were generated by transducing each mutant strain with a phage lysate produced in the *hly::himar1* background, as previously described (16). Briefly, U153 phage were mixed with the appropriate donor strain and incubated at 30 °C in LB soft agar overnight. Phage lysates were eluted from agar, filter-sterilized, and added to recipient strains for 30 minutes at room temperature. Transductions were plated on antibiotic containing agar and incubated at 37 °C. Insertions in *hly* were confirmed by PCR.

### Growth curves and peroxide toxicity

For anaerobic growth, colonies were inoculated into TSB and incubated at 37°C in closed containers containing anaerobic gas-generating pouches (GasPak EZ; BD). Anaerobic overnight cultures were normalized to OD_600_ 0.02 in 25 mL of TSB in 250 mL flasks and grown aerobically with shaking at 37°C. At each time point, bacteria were serially diluted, plated on BHI agar, and grown anaerobically to enumerate CFU.

To assess the response to acute peroxide toxicity, bacteria grown overnight in TSB at 37°C, shaking were back-diluted to OD_600_ = 0.1 and incubated for 2 hours to OD_600_ ~ 0.6. Hydrogen peroxide (120 mM, Sigma) was added and cultures were incubated with shaking for 1 hour. At each time point, bacterial cultures were serially diluted and plated on BHI to enumerate CFU.

### Quantitative RT-PCR of bacterial transcripts

Bacteria grown anaerobically overnight in degassed TSB were normalized to OD_600_ = 0.02 in 50 mL media in 500 mL flasks and incubated at 37°C in the dark, with shaking. At each time point, 5-7 mL of bacterial culture were removed, mixed with ice-cold methanol (1:1), pelleted by centrifugation, flash frozen, and stored at −80°C. Nucleic acids were harvested as previously described using phenol:chloroform extraction and bead beating (16). RNA was precipitated overnight, washed in ethanol, and RT-PCR was performed with iScript Reverse Transcriptase (Bio-Rad). Quantitative PCR was performed with iTaq Universal SYBR Green Supermix (Bio-Rad) according to manufacturers’ recommendations. Transcripts were normalized to that of 16S rRNA and fold-change was calculated using the comparative C_T_ method.

### Macrophage growth curves

BMDMs were seeded at a concentration of 6 × 10^5^ cells per well in TC-treated 24-well plates the day before infection. BMDMs were activated by incubating the monolayer with recombinant murine IFNγ (100 ng/mL, PeproTech) overnight and during infection. Overnight bacterial cultures were grown anaerobically in TSB (Δ*fri* aerobically) at 30°C statically, washed twice with PBS, and resuspended in warmed BMDM media (16). BMDMs were infected at an MOI of 0.1 for 30 minutes before cells were washed twice with PBS and then BMDM media containing gentamicin (50 μg/mL) was added to each well. To measure bacterial growth, cells were washed twice with PBS and then lysed by incubating with 250 μL cold PBS with 0.1% Triton X-100 for 5 min at room temperature, followed by serial dilutions and plating on BHI agar to enumerate CFU. For the *phox*^-/-^ growth curves all strains were incubated aerobically in TSB.

### Infections and flow cytometry

J774 cells were plated in 12-well TC-treated dishes at 10^6^ cells per well. Generally, *L. monocytogenes* transcriptional reporter strains were grown to mid-log phase at 37°C with shaking. After being washed twice and resuspended in PBS, bacterial suspensions were added to the cells at an MOI=10. One hour post-infection, cells were washed twice with PBS and media containing gentamicin (50 μg/mL) was added to each well. 6 hours post-infection the cells were washed twice with PBS, treated with 0.25% trypsin (Gibco), and resuspended in an equal volume of media. The cells were then fixed with 2% formaldehyde, washed twice with flow buffer (PBS containing 5% FBS), and resuspended in 300 μL flow buffer. Flow cytometry was performed on an LSRII flow cytometer (BD) and analyzed using FlowJo (FlowJo, LLC). Cells were discriminated from debris by forward scatter area (FSC-A) and side scatter area (SSC-A). Single cells were gated using FSC-A and forward scatter height (FSC-H). The GFP gate was set to include 5% of the uninfected sample; this gate represents infected cells, and is referred to as GFP-positive. Within the infected cell population, the RFP gate was set to include 5% of cells infected with pH-gfp; this gate is referred to as RFP-positive. Reported values represent %RFP-positive cells within GFP-positive cells of a given sample.

### Plaque Assays

Plaque assays were performed as previously described (16, 27, 54, 55). Briefly, TC-treated 6-well dishes were seeded with 1.2 x 10^6^ L2 murine fibroblasts per well. *L. monocytogenes* strains were incubated in BHI overnight at 30°C, static. Overnight cultures were diluted 1:10 in sterile PBS, and 5-10 μL was used to infect each well. One hour post-infection, cells were washed twice with PBS and 3 mL of molten agarose-DMEM solution was added to each well. This solution consisted of a 1:1 mixture of 2X DMEM (Gibco) and 1.4% SuperPure agarose LE (U.S. Biotech Sources, LLC) containing gentamicin (10 μg/mL). Three days post-infection, 2 mL of molten agarose-DMEM solution containing neutral red was added to each well to visualize plaques. After 24 hours, the plates were scanned, and the plaque areas measured using ImageJ software (56). The area of at least 20 plaques was measured for each strain and normalized to that of wt.

## ACKNOWLEDGMENTS

The authors would like to thank Steve Libby and Ferric Fang (UW) for the *phox*^-/-^ mice and Savannah Bertolli and the Mougous Lab (UW) for technical assistance and time in the anaerobic chamber. Research in the Reniere Lab is funded by NIH R01 AI132356. M.R.C. was funded by the Howard Hughes Medical Institute through the James H. Gilliam Fellowships for Advanced Study program (GT11030). C.R.H. was supported by NIH grant T32AI055396. The funders had no role in study design, data collection and interpretation, or the decision to submit the work for publication. The authors do not have a conflict of interest to declare.

## REFERENCES

1. Reniere ML. 2018. Reduce, Induce, Thrive: Bacterial Redox Sensing during Pathogenesis. Journal of Bacteriology 200:903.

2. Nauseef WM, Clark RA. 2019. Intersecting Stories of the Phagocyte NADPH Oxidase and Chronic Granulomatous Disease, p. 3–16. In Knaus, UG, Leto, TL (eds.), NADPH Oxidases: Methods and Protocols. Springer, New York, NY.

3. Dinauer MC. 2005. Chronic granulomatous disease and other disorders of phagocyte function. Hematology Am Soc Hematol Educ Program 2005:89–95.

4. van den Berg JM, van Koppen E, Ahlin A, Belohradsky BH, Bernatowska E, Corbeel L, Español T, Fischer A, Kurenko-Deptuch M, Mouy R, Petropoulou T, Roesler J, Seger R, Stasia M-J, Valerius NH, Weening RS, Wolach B, Roos D, Kuijpers TW. 2009. Chronic granulomatous disease: the European experience. PLoS ONE 4:e5234.

5. Imlay JA. 2008. Cellular defenses against superoxide and hydrogen peroxide. Annu Rev Biochem 77:755–776.

6. Mishra S, Imlay JA. 2012. Why do bacteria use so many enzymes to scavenge hydrogen peroxide? Archives of Biochemistry and Biophysics 525:145–160.

7. Park S, You X, Imlay JA. 2005. Substantial DNA damage from submicromolar intracellular hydrogen peroxide detected in Hpx-mutants of Escherichia coli. Proc Natl Acad Sci USA 102:9317–9322.

8. Ruhland BR, Reniere ML. 2018. Sense and sensor ability: redox-responsive regulators in Listeria monocytogenes. Curr Opin Microbiol 47:20–25.

9. Freitag NE, Port GC, Miner MD. 2009. Listeria monocytogenes - from saprophyte to intracellular pathogen. Nat Rev Microbiol 7:623–628.

10. Reniere ML, Whiteley AT, Hamilton KL, John SM, Lauer P, Brennan RG, Portnoy DA. 2015. Glutathione activates virulence gene expression of an intracellular pathogen. Nature 517:170–173.

11. Mains DR, Eallonardo SJ, Freitag NE. 2021. Identification of Listeria monocytogenes genes contributing to oxidative stress resistance under conditions relevant to host infection. Infection and Immunity https://doi.org/10.1128/IAI.00700-20.

12. Cesinger MR, Thomason MK, Edrozo MB, Halsey CR, Reniere ML. 2020. Listeria monocytogenes SpxA1 is a global regulator required to activate genes encoding catalase and heme biosynthesis enzymes for aerobic growth. Mol Microbiol https://doi.org/10.1111/mmi.14508.

13. Savelli B, Li Q, Webber M, Jemmat AM, Robitaille A, Zamocky M, Mathé C, Dunand C. 2019. RedoxiBase: A database for ROS homeostasis regulated proteins. Redox Biology 26:101247.

14. Olsen KN, Larsen MH, Gahan CGM, Kallipolitis B, Wolf XA, Rea R, Hill C, Ingmer H. 2005. The Dps-like protein Fri of Listeria monocytogenes promotes stress tolerance and intracellular multiplication in macrophage-like cells. Microbiology (Reading, Engl) 151:925–933.

15. Dussurget O, Dumas E, Archambaud C, Chafsey I, Chambon C, Hébraud M, Cossart P. 2005. Listeria monocytogenes ferritin protects against multiple stresses and is required for virulence. FEMS Microbiol Lett 250:253–261.

16. Reniere ML, Whiteley AT, Portnoy DA. 2016. An In Vivo Selection Identifies Listeria monocytogenes Genes Required to Sense the Intracellular Environment and Activate Virulence Factor Expression. PLoS Pathog 12:e1005741.

17. Cepeda JA, Millar M, Sheridan EA, Warwick S, Raftery M, Bean DC, Wareham DW. 2006. Listeriosis due to infection with a catalase-negative strain of Listeria monocytogenes. J Clin Microbiol 44:1917–1918.

18. Leblond-Francillard M, Gaillard JL, Berche P. 1989. Loss of catalase activity in Tn1545-induced mutants does not reduce growth of Listeria monocytogenes in vivo. Infect Immun 57:2569–2573.

19. Dons LE, Mosa A, Rottenberg ME, Rosenkrantz JT, Kristensson K, Olsen JE. 2014. Role of the Listeria monocytogenes 2-Cys peroxiredoxin homologue in protection against oxidative and nitrosative stress and in virulence. Pathog Dis 70:70–74.

20. Kim K-P, Hahm B-K, Bhunia AK. 2007. The 2-cys peroxiredoxin-deficient Listeria monocytogenes displays impaired growth and survival in the presence of hydrogen peroxide in vitro but not in mouse organs. Curr Microbiol 54:382–387.

21. Milazzo L, Hofbauer S, Howes BD, Gabler T, Furtmüller PG, Obinger C, Smulevich G. 2018. Insights into the Active Site of Coproheme Decarboxylase from Listeria monocytogenes. Biochemistry 57:2044–2057.

22. Cosgrove K, Coutts G, Jonsson I-M, Tarkowski A, Kokai-Kun JF, Mond JJ, Foster SJ. 2007. Catalase (KatA) and Alkyl Hydroperoxide Reductase (AhpC) Have Compensatory Roles in Peroxide Stress Resistance and Are Required for Survival, Persistence, and Nasal Colonization in Staphylococcus aureus. Journal of Bacteriology 189:1025–1035.

23. Lauer P, Chow MYN, Loessner MJ, Portnoy DA, Calendar R. 2002. Construction, characterization, and use of two Listeria monocytogenes site-specific phage integration vectors. 184:4177–4186.

24. Collins B, Curtis N, Cotter PD, Hill C, Ross RP. 2010. The ABC transporter AnrAB contributes to the innate resistance of Listeria monocytogenes to nisin, bacitracin, and various beta-lactam antibiotics. Antimicrobial Agents and Chemotherapy 54:4416–4423.

25. Torres VJ, Stauff DL, Pishchany G, Bezbradica JS, Gordy LE, Iturregui J, Anderson KL, Dunman PM, Joyce S, Skaar EP. 2007. A Staphylococcus aureus regulatory system that responds to host heme and modulates virulence. Cell Host Microbe 1:109–119.

26. Herskovits AA, Auerbuch V, Portnoy DA. 2007. Bacterial ligands generated in a phagosome are targets of the cytosolic innate immune system. PLoS Pathog 3:e51.

27. Sun AN, Camilli A, Portnoy DA. 1990. Isolation of Listeria monocytogenes small-plaque mutants defective for intracellular growth and cell-to-cell spread. Infect Immun 58:3770–3778.

28. Bsat N, Chen L, Helmann JD. 1996. Mutation of the Bacillus subtilis alkyl hydroperoxide reductase (ahpCF) operon reveals compensatory interactions among hydrogen peroxide stress genes. Journal of Bacteriology 178:6579–6586.

29. Chen L, Keramati L, Helmann JD. 1995. Coordinate regulation of Bacillus subtilis peroxide stress genes by hydrogen peroxide and metal ions. PNAS 92:8190–8194.

30. Seaver LC, Imlay JA. 2001. Alkyl hydroperoxide reductase is the primary scavenger of endogenous hydrogen peroxide in Escherichia coli. Journal of Bacteriology 183:7173–7181.

31. Broden NJ, Flury S, King AN, Schroeder BW, Coe GD, Faulkner MJ. 2016. Insights into the Function of a Second, Nonclassical Ahp Peroxidase, AhpA, in Oxidative Stress Resistance in Bacillus subtilis. Journal of Bacteriology 198:1044–1057.

32. Su M, Cavallo S, Stefanini S, Chiancone E, Chasteen ND. 2005. The So-Called Listeria innocua Ferritin Is a Dps Protein. Iron Incorporation, Detoxification, and DNA Protection Properties. Biochemistry 44:5572–5578.

33. Bozzi M, Mignogna G, Stefanini S, Barra D, Longhi C, Valenti P, Chiancone E. 1997. A novel non-heme iron-binding ferritin related to the DNA-binding proteins of the Dps family in Listeria innocua. J Biol Chem 272:3259–3265.

34. Hébraud M, Guzzo J. 2000. The main cold shock protein of Listeria monocytogenes belongs to the family of ferritin-like proteins. FEMS Microbiology Letters 190:29–34.

35. Mohamed W, Darji A, Domann E, Chiancone E, Chakraborty T. 2006. The ferritin-like protein Frm is a target for the humoral immune response to Listeria monocytogenes and is required for efficient bacterial survival. Mol Genet Genomics 275:344–353.

36. Mohamed W, Sethi S, Darji A, Mraheil MA, Hain T, Chakraborty T. 2010. Antibody Targeting the Ferritin-Like Protein Controls Listeria Infection. Infection and Immunity 78:3306–3314.

37. Polidoro M, De Biase D, Montagnini B, Guarrera L, Cavallo S, Valenti P, Stefanini S, Chiancone E. 2002. The expression of the dodecameric ferritin in Listeria spp. is induced by iron limitation and stationary growth phase. Gene 296:121–128.

38. Proctor RA, von Eiff C, Kahl BC, Becker K, McNamara P, Herrmann M, Peters G. 2006. Small colony variants: a pathogenic form of bacteria that facilitates persistent and recurrent infections. Nat Rev Microbiol 4:295–305.

39. Hammer ND, Reniere ML, Cassat JE, Zhang Y, Hirsch AO, Indriati Hood M, Skaar EP. 2013. Two heme-dependent terminal oxidases power Staphylococcus aureus organ-specific colonization of the vertebrate host. MBio 4:e00241–13.

40. Corbett D, Goldrick M, Fernandes VE, Davidge K, Poole RK, Andrew PW, Cavet J, Roberts IS. 2017. Listeria monocytogenes has both a bd-type and an aa3-type terminal oxidase which allow growth in different oxygen levels and both are important in infection. Infect Immun 85:e00354–17.

41. Choby JE, Skaar EP. 2016. Heme Synthesis and Acquisition in Bacterial Pathogens. J Mol Biol 428:3408–3428.

42. Jongbloed JDH, Grieger U, Antelmann H, Hecker M, Nijland R, Bron S, Dijl JMV. 2004. Two minimal Tat translocases in Bacillus. Molecular Microbiology 54:1319–1325.

43. Miethke M, Monteferrante CG, Marahiel MA, van Dijl JM. 2013. The Bacillus subtilis EfeUOB transporter is essential for high-affinity acquisition of ferrous and ferric iron. Biochimica et Biophysica Acta (BBA) - Molecular Cell Research 1833:2267–2278.

44. Ledala N, Sengupta M, Muthaiyan A, Wilkinson BJ, Jayaswal RK. 2010. Transcriptomic response of Listeria monocytogenes to iron limitation and Fur mutation. Appl Environ Microbiol 76:406–416.

45. dos Santos PT, Larsen PT, Menendez-Gil P, Lillebæk EMS, Kallipolitis BH. 2018. Listeria monocytogenes Relies on the Heme-Regulated Transporter hrtAB to Resist Heme Toxicity and Uses Heme as a Signal to Induce Transcription of lmo1634, Encoding Listeria Adhesion Protein. Front Microbiol 9.

46. Hingston PA, Piercey MJ, Hansen LT. 2015. Genes Associated with Desiccation and Osmotic Stress in Listeria monocytogenes as Revealed by Insertional Mutagenesis. Appl Environ Microbiol 81:5350–5362.

47. Nakano S, Küster-Schöck E, Grossman AD, Zuber P. 2003. Spx-dependent global transcriptional control is induced by thiol-specific oxidative stress in Bacillus subtilis. Proc Natl Acad Sci USA 100:13603–13608.

48. Vasconcelos JA, Deneer HG. 1994. Expression of superoxide dismutase in Listeria monocytogenes. Appl Environ Microbiol 60:2360–2366.

49. Archambaud C, Nahori M-A, Pizarro-Cerda J, Cossart P, Dussurget O. 2006. Control of Listeria superoxide dismutase by phosphorylation. J Biol Chem 281:31812–31822.

50. Sauer J-D, Sotelo-Troha K, von Moltke J, Monroe KM, Rae CS, Brubaker SW, Hyodo M, Hayakawa Y, Woodward JJ, Portnoy DA, Vance RE. 2011. The N-ethyl-N-nitrosourea-induced Goldenticket mouse mutant reveals an essential function of Sting in the in vivo interferon response to Listeria monocytogenes and cyclic dinucleotides. Infect Immun 79:688–694.

51. Bishop DK, Hinrichs DJ. 1987. Adoptive transfer of immunity to Listeria monocytogenes. The influence of in vitro stimulation on lymphocyte subset requirements. J Immunol 139:2005–2009.

52. Bécavin C, Bouchier C, Lechat P, Archambaud C, Creno S, Gouin E, Wu Z, Kühbacher A, Brisse S, Pucciarelli MG, García-Del Portillo F, Hain T, Portnoy DA, Chakraborty T, Lecuit M, Pizarro-Cerda J, Moszer I, Bierne H, Cossart P. 2014. Comparison of Widely Used Listeria monocytogenes Strains EGD, 10403S, and EGD-e Highlights Genomic Variations Underlying Differences in Pathogenicity. MBio 5.

53. Simon R, Priefer U, Pühler A. 1983. A broad host range mobilization system for in vivo genetic engineering: transposon mutagenesis in Gram-negative bacteria. Nat Biotechnol 784–791.

54. Whiteley AT, Ruhland BR, Edrozo MB, Reniere ML. 2017. A Redox-Responsive Transcription Factor Is Critical for Pathogenesis and Aerobic Growth of Listeria monocytogenes. Infect Immun 85:e00978–16.

55. Ruhland BR, Reniere ML. 2020. YjbH Requires Its Thioredoxin A for the Nitrosative Stress Response, Cell-to-Cell Spread, and Protein-Protein Interactions in *Listeria monocytogenes*. J Bacteriol 202:e00099–20, http://jb/202/12/JB.00099-20.atom.

56. Schneider CA, Rasband WS, Eliceiri KW. 2012. NIH Image to ImageJ: 25 years of image analysis. Nat Methods 9:671–675.

